# Organisation of the nervous system in cysts of the freshwater tardigrade *Thulinius ruffoi* (Parachela, Isohypsibioidea: Doryphoribiidae)

**DOI:** 10.1101/2023.08.14.553198

**Authors:** Kamil Janelt, Izabela Poprawa

## Abstract

Encystment is a natural process that involves cyst formation, and at least some species of tardigrades can form cysts. However, the encystment process and cyst structure among tardigrades are still poorly understood. Despite some aspects of the encysted animals’ system organisation being examined in the past, the morphology and structure of the nervous system have never been thoroughly investigated. The present study covers anatomical, histological and morphological details and proposes physiological aspects of the nervous system in encysted *Thulinius ruffoi* up to 11 months duration in encystment. This is the first record of the nervous system organisation in a species belonging to the family Doryphoribiidae and describes morphological changes that occur during cyst formation.

## 1. INTRODUCTION

The phylum Tardigrada, colloquially known as water bears, is a commonly occurring group of micrometazoan invertebrates. Traditionally, three classes are distinguished among the Tardigrada: Heterotardigrada, Eutardigrada, and Mesotardigrada. However, the class Mesotardigrada currently has a status of *nomen dubium*. Heterotardigrada is divided into the orders Arthrotardigrada and Echiniscoidea, whereas Eutardigrada consists of the orders Apochela and Parachela (Nelson et al., 2015; Degma & Guidetti, 2023). The general morphology of tardigrades includes a segmented body plan in which the head and four body segments are present, with a pair of legs characterising each of the trunk segments (Ramazzotti & Maucci, 1983; Møbjerg et al., 2018). Segmentation is also well documented in the organisation of the nervous system of these animals, with the brain and ganglionated nervous system of the trunk representing the main part. Also, all innervations, including leg and peripheral ganglia are a part of this system (Ramazzotti & Maucci, 1983; Møbjerg et al., 2018; Mayer et al., 2013a, b). Sensory organs in Heterotardigrada are characterised by numerous externally protruded sensory appendages on the head (cirri and clavae) and trunk (cirri) and also occur on the legs (Møbjerg et al., 2018). Although the appendages are mostly paired, unpaired sensory organs on the head and the trunk may be present (Degma & Guidetti, 2018; Møbjerg et al., 2018). In the dubious Mesotardigrada (Grothman et al., 2017), eye spots, peribuccal papillae and lateral cirri were described in the only representative species, *Thermozodium esakii* (Rahm, 1937). Eye spots are also present in some representatives of both heterotardigrades and eutardigrades (Kristensen, 1982; Ramazzotti & Maucci, 1983; Greven, 2007; Degma & Guidetti, 2018; Møbjerg et al., 2018). Among eutardigrades, the sensory structures are much more reduced externally, except the mouth, which is surrounded by sense organs called peribuccal (circumoral) and infrabuccal structures (Møbjerg et al., 2018). The cephalic region of eutardigrades is characterised by the presence of sensory fields and foregut-associated sense organs (Walz, 1978; Wiederhöft & Greven, 1996, 1999; Zantke et al., 2008; Biserova & Kuznetsova, 2012; Persson et al., 2012; Mayer et al., 2013a, b; Gross & Mayer, 2015; Gross et al., 2021). Some extracephalic sensory fields, such as the cloacal sensory field and a putative sensory field associated with the flat bulge on the hind legs, were proposed in *Macrobiotus kyoukenus* by Cesari et al. (2022).

In 1906, histological methods were used for the first time to analyse the internal organisation of tardigrades (Basse, 1906), and the importance of the future anatomical and histological studies of individuals during encystment was noted (Lauterborn, 1906). The first observations on sectioned cysts of *Macrobiotus lacustris*, presently considered to be *Thulinius augusti* (according to Degma & Guidetti, 2023), revealed valuable information on the internal organisation of encysted animals (von Wenck, 1914), including the presence of eye spots (also mentioned in encysted animals e.g. by Murray, 1907a, b; Hansen & Katholm, 2002; Guidetti et al., 2006). Unfortunately, von Wenck’s study did not provide much information about the nervous system, and there were no further histological studies until recently.

Previously, the presence of peribuccal structures in encysted *Thulinius ruffoi* dissected from the cuticular capsule was reported, as well as the general morphology of the cysts and some aspects of behaviour related to cyst formation and excystment (Janelt & Poprawa, 2020). In the present study, based on the importance of histological studies, the organisation of the nervous system of *Thu. ruffoi* was analysed by examining sectioned cysts. Moreover, morphological changes during encystment were also examined and are described here.

## 2. MATERIAL AND METHODS

### 2.1. Specimens and culturing

Individuals of *Thu*. *ruffoi* (Thu.ruf_PL.014 strain) used in this study came from the samples collected from a wastewater treatment plant as described by Sobczyk et al. (2015) and kindly provided by the Michalczyk Lab (Jagiellonian University, Kraków, Poland) in 2014. The specimens were cultured at 19 °C in plastic Petri dishes and 24-well scratched polystyrene plates with a culture medium composed of distilled water and Żywiec Zdrój spring water (1:1) and fed with a mixture of *Chlorella* sp. and *Chlorococcum* sp. *ad libitum* as described (Janelt et al., 2019, 2020; Janelt & Poprawa, 2020). Observations were performed at RT (about 25 °C). During the observations, cysts were not found.

#### 2.1.1. Obtaining cysts

Cysts for this research were obtained under artificial laboratory conditions (Figure S1). Based on the protocol in a previous paper (Janelt & Poprawa, 2020), each plate dedicated to cyst procurement was transferred to a temperature of 6.5 °C. Observations were performed at RT for the shortest possible time. After that, the plates were transferred again to 6.5 °C.

The first set of plates (five scratched and two unscratched) was designed to obtain cysts with a well-established time of encystment duration (cysts with a known ’’age’’ were defined as the time elapsed from cyst formation to its isolation). Only one active, randomly selected individual was placed on the scratched bottom of each well. Then each plate was checked to confirm the presence of one specimen in each well. If all animals were present, each well was filled with 1 ml of medium, and a mixture of algae was added.

While under observation, all deposited eggs and/or hatched individuals in the five scratched plates were removed from the wells, leaving only the initially placed animals. Each animal on these plates was observed every 24–48 hours until the day of cyst formation. Six freshly formed cysts (up to 2 days old) were removed, cleaned gently by washing each cyst several times with a fresh portion of distilled water and then fixed in 2.5% glutaraldehyde for further analyses (Table S1). Other freshly formed cysts were removed, cleaned gently and placed in a drop of fresh distilled water on two unscratched 24-well polystyrene plates. After the cyst was placed, the well was checked to confirm the presence of the cyst. The well was then filled with 1 ml of fresh distilled water, and food was not added. The date the cyst was obtained and the time to the end of incubation were noted on the plate cover over each well. The prepared plates were transferred to 6.5 °C for incubation until the specified time – 1, 3, 6, 8-9 or 11 mths. The plates were checked periodically with each new cyst added. After a specific incubation time, another thirteen cysts were recovered, cleaned, and fixed using 2.5% glutaraldehyde (Table S1).

The second set of plates (four scratched plates) was used to obtain cysts without an established time of encystment duration. After placing one non-encysted animal in a drop of distilled water on a scratched bottom of the well, the plates were filled with medium and food was added. The animals on these plates reproduced and grew. However, neither eggs nor hatched animals were removed. The wells of the plates were occasionally observed to check if any cysts were present (the min/max interval between checks was seven days and six months, respectively). Less frequent observations did not permit estimating a more precise time when the cyst was formed. In addition, it was not always possible to tell whether the originally placed individual or another had formed a cyst. Eleven cysts were isolated from the plates, cleaned gently, and fixed in 2.5% glutaraldehyde for further analysis (Table S1).

### 2.2. Sectioning cysts

#### 2.2.1. LM, TEM, STEM

Twenty-nine of thirty isolated cysts (Table S1) were washed a few times in fresh distilled water to clean them. Afterwards, they were fixed in 2.5% glutaraldehyde in 0.1 M phosphate buffer (pH of 7.4) at 4 °C for at least 2 hours in Eppendorf tubes. If necessary, the glutaraldehyde retention period was extended to 6 days to collect more material for further processing steps. Then, the material was washed in phosphate buffer for 1.5 h at RT or overnight at 4 °C, depending on the time in glutaraldehyde. After that, the material was processed according to the previously described methodology (Janelt & Poprawa, 2020). The material was embedded in epoxy resin using an Epoxy Embedding Medium Kit (Sigma-Aldrich).

Semi [and ultra[thin sections were cut on a Leica EM UC7 ultramicrotome. Semi-thin sections of 250–500 nm thickness were mounted in a drop of distilled water on a microscope slide, dried and stained with 1% methylene blue in 1% borax. After staining, the slides were washed with distilled water, dried and then DPX mounting medium and a coverslip were added. Ultra[thin sections of 50–70 nm thickness were mounted on copper grids (Agar Scientific, UK). Some of them were coated with formvar. Sections on the grids were counterstained with uranyl acetate and lead citrate.

#### 2.2.2. SBEM

One six-month-old cyst of *Thu*. *ruffoi* (Table S1) was used to acquire serial EM images with a serial block-face scanning electron microscope (SBEM). The cyst was prepared as described previously (Janelt et al., 2020), in general according to Deerinck et al. (2010).

The succeeding steps were similar to those used for TEM, including incubation of the cyst in the acetone-resin mixture and subsequent evaporation of the acetone. Each cyst was embedded in epoxy resin with single-walled carbon nanotubes (US Research Nanomaterials), sandwiched between ACLAR (Ted Pella) sheets, and left to polymerise at 60 °C for five days. The cyst was cut from the larger amount of the epoxy resin after removal of the ACLAR layers and embedded on an aluminium pin (metal rivets, Oxford Instruments) using two-component transparent epoxy glue mixed with carbon nanotubes. The pin was left to dry, after which a portion of epoxy resin with a sample on the pin’s head was trimmed to remove excess resin. Next, the sample was coated with silver paint (Ted Pella) and dried for 24 h.

The cyst was cut into serial sections of 150 nm thickness using a Zeiss Sigma VP Scanning Electron Microscope equipped with a backscatter electron detector and ultramicrotome module 3View2 (Gatan). The surface was scanned before cutting each section off (scanning parameters: variable pressure 25 Pa, EHT 3 kV, aperture 30 µm, dwell time 3 µs), producing images mostly at 2500 x magnification (image parameters: pixel size 16.8 nm, 5632 x 4608 px). At depths where the cyst extended beyond the scan area, scans at lower magnifications of 1800, 2000 or 2300 x were made to capture the whole area. In these cases, such scans were taken every ten sections (5632 x 4608 px with a pixel size of 23.3 nm, 21.0 nm and 18.2 nm, respectively, depending on the magnification).

### 2.3. Three-dimensional visualisation

Three-dimensional visualisations were performed on a set of scans from the 6-month-old cyst of *Thu*. *ruffoi*. Scans of the sections were aligned manually using landmarks, after which each modelling element was manually defined. Alignment and segmentation were performed using the TrackEM plugin. Any losses of information caused by biological material beyond the scanning area were supplemented by interpolation. The gaps were filled based on the same regions above and below each gap, which were captured on sections made at lower magnifications. Then the 3D Viewer plugin was used for three-dimensional rendering. Both plugins are a part of Fiji, a scientific image processing package based on ImageJ (Schindelin et al., 2012), which is open-source software (https://fiji.sc/).

### 2.4. Data acquisition and image processing

All observations of living cultures were performed at room temperature (RT) using an Olympus SZ40 stereoscopic microscope equipped with a black-and-white stage plate and an Olympus Highlight 2100 microscopy illuminator. Data from the slides were collected the next day after mounting the specimens with DPX medium and a coverslip. Data were collected using the Olympus BX60 microscope equipped with an Olympus XC50 microscopic camera and appropriate Olympus software (CellSense).

The grids with ultrathin sections were analysed with a Hitachi H500 transmission electron microscope at 75 kV. Documentation from the sections was captured on photographic film (Carestream Electron microscope Film 4489, Sigma-Aldrich), which was developed and scanned into an electronic version. Some grids with the counterstained sections were also analysed using an ultra-high resolution scanning electron microscope (Hitachi UHR FE-SEM SU 8010) equipped with a TE detector to examine the sections at a low voltage of 25 kV. Digital data were collected from the Hitachi UHR FE-SEM SU 8010 and the Zeiss Sigma VP Scanning Electron Microscope with the appropriate device software.

All the mentioned microscopes and other equipment, except the Zeiss Sigma VP Scanning Electron Microscope equipped with an ultramicrotome module (which is available at the Nencki Institute of Experimental Biology Polish Academy of Sciences; Warszawa, Poland), and the CorelDraw Graphics Suite used for the final assembly of the figures are available at the Institute of Biology, Biotechnology and Environmental Protection (Faculty of Natural Sciences, University of Silesia in Katowice).

### 2.5. Taxonomic nomenclature

All abbreviations of the generic names were consistent with the International Code of Zoological Nomenclature (ICZN), according to the recommendations of Perry et al. (2019, 2021). Other taxonomic and systematic information was from the current (42nd edition) checklist of Tardigrada species (Degma & Guidetti, 2023).

## 3. RESULTS

### 3.1. Cephalic nervous system

Analysis of the neural tissue within the cephalic region (Figures 1, 2 and 3) showed the presence of dorsolateral (DLc), ventral (Vc), paired anterolateral (ALc) and paired posterolateral (PLc) neural clusters. The reconstructed neural cephalic structures exhibited bilateral symmetry (Figure 1a–i, Video S1). In the cephalic neural clusters, surrounded externally by a basal lamina, were regions rich in neurites and somata-rich regions containing the nerve cell bodies. The fibre regions had a multi-membranous appearance and showed the boundaries between projections of the neurons. Synapses were observed in the sections from the cephalic neural tissue (Figure 1h–l).

**Figure 1.**
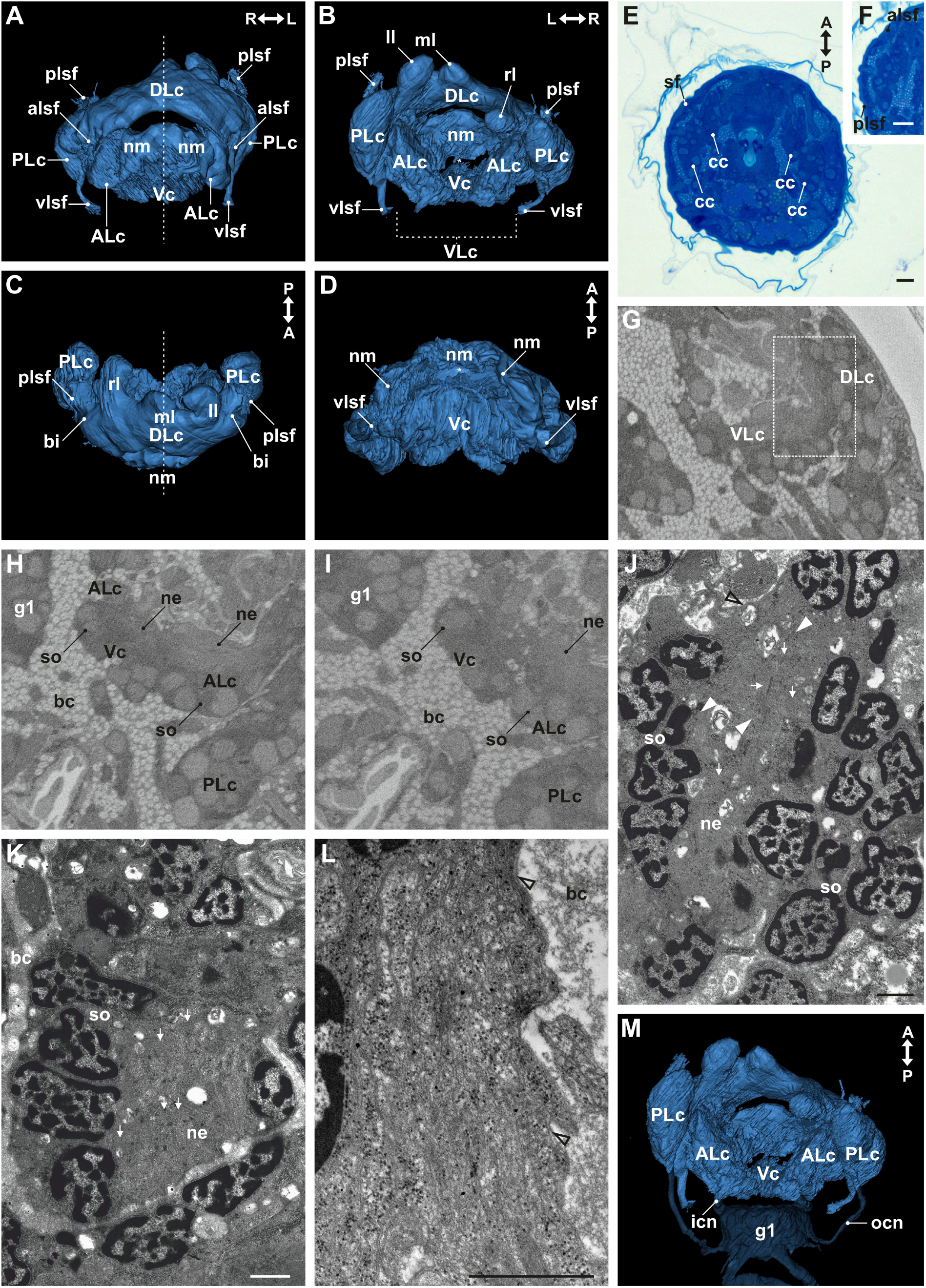
Cephalic neural structures of encysted *Thu*. *ruffoi*. (A–L) cephalic neural structures and (M) relation between them and the trunk nervous system. (A–D, L) cephalic neural structures of a 6-month-old cyst; (E, F) 11-month-old cyst; (G–I) sections from the same specimen as (A–D, L); (J–L) Fragments of the cephalic clusters from three different encysted animals at the unknown time of encystment duration. (A–D, L) 3D visualisation based on SBEM images; (E, F) LM, scale bar 10 µm; (G–I) SBEM; (J, L) STEM and (K) TEM, scale bar 1 µm. Note that L and R denote the left and right sides while A and P refer to the anteroposterior axis. Anterolateral cluster (ALc); anterolateral sensory field (alsf); body cavity (bc); bridge (bi); cephalic nervous cluster (cc); dorsolateral cluster (DLc); ventral ganglion (g); inner connective (icn); left lobe (ll); medial lobe (ml); neurites (ne); protruding extensions of the VLc (nm); outer connective (ocn); posterolateral nervous cluster (PLc); posterolateral sensory field (plsf); right lobe (rl); sensory field (sf); somata (so); ventral nervous cluster (Vc); ventrolateral complex (VLc); ventrolateral sensory field (vlsf); basal lamina (empty arrowhead); interspaces between neural cells (filled arrowhead); a slit through which the mouth area retracts (asterisk); synapse with synaptic vesicles (thin arrows). Note that the dotted rectangle indicates the connection between protruding extensions of the VLc and the dorsolateral cluster.

**Figure 2.**
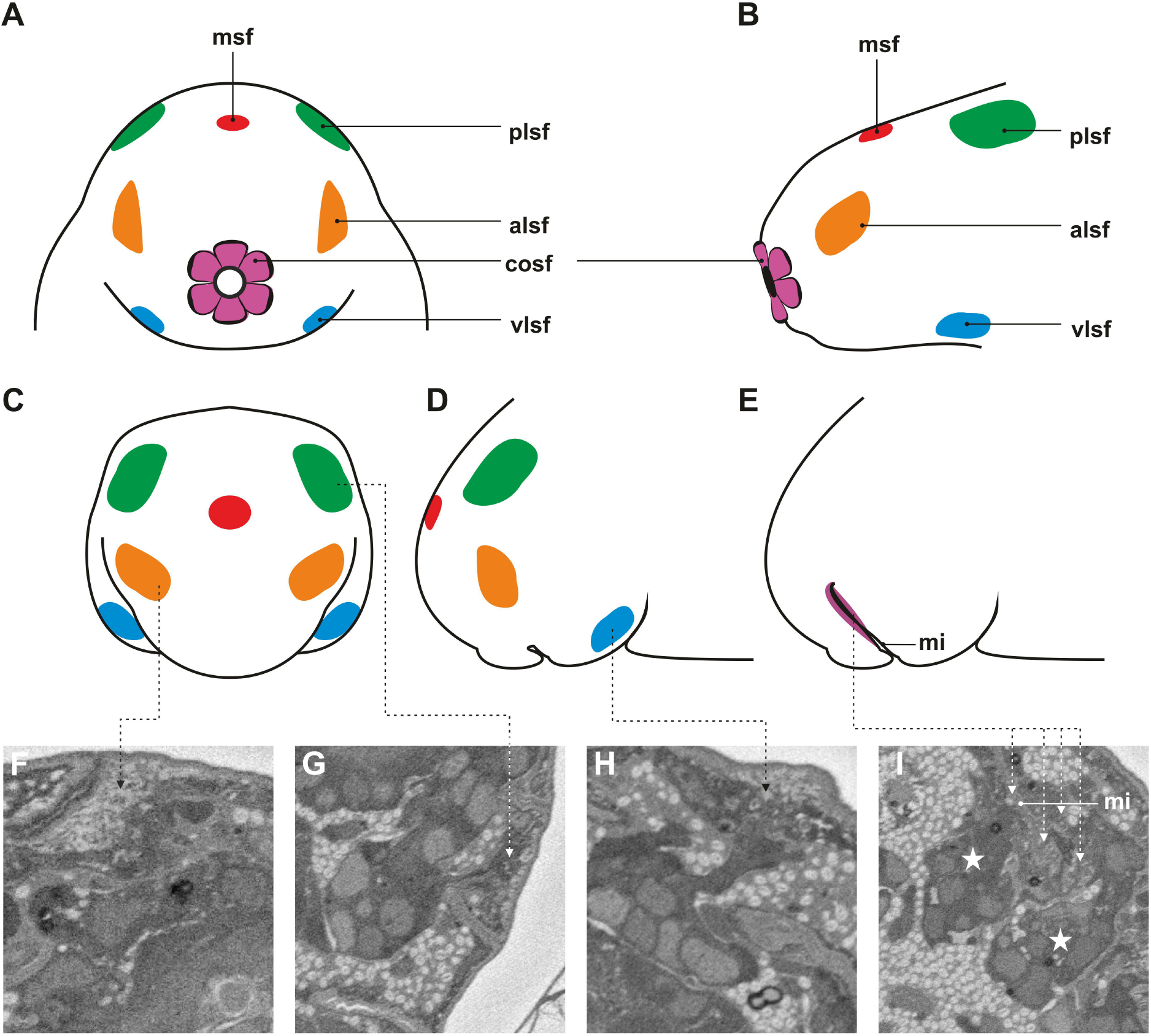
Integument-associated sensory fields of *Thu*. *ruffoi*. (A, B) non-encysted animal; (C–I) encysted animal. (A, B) external integument-associated sensory fields in non-encysted animals; (C, D) external sensory fields in encysted animals; (A, C) frontal view; (B, D) lateral view; (E) schematic of inwards retraction of the mouth region in lateral view; (F–I) integument-associated sensory fields, SBEM; (F) anterolateral sensory field; (G) posterolateral sensory field; (H) ventrolateral sensory field; (I) circumoral sensory field. Note the changed position of the sensory fields. The presence of the medial sensory field is hypothesised due to lack of data from this region. Anterolateral sensory field (alsf, orange); circumoral sensory field (cosf, violet); mouth area invagination (mi); median sensory field (msf, red); posterolateral sensory field (plsf, green); elements forming the circumbuccal ring-shaped structure (star) innervating the mouth region; ventrolateral sensory field (vlsf, blue). The median sensory field is probably present, but its occurrence could not be verified in the present study.

**Figure 3.**
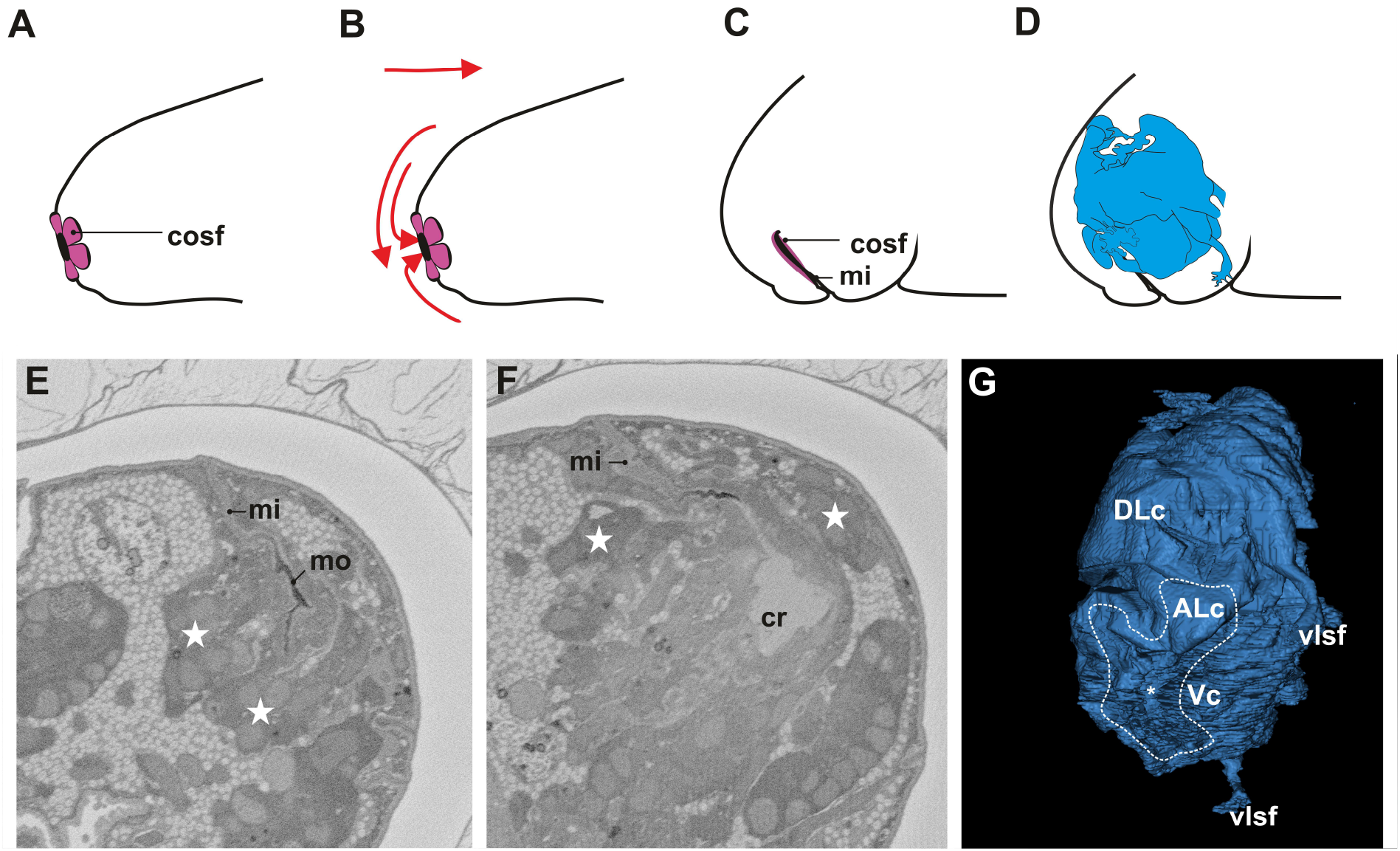
Cephalic region of *Thu*. *ruffoi* and changes during cyst formation. (A, B) non-encysted animal; (B) schematic of changes (red arrows) during which the retraction of the mouth area, the tilt of the head towards the ventral side, and contraction of the head into the rest of the body occur during cyst formation; (C, D) encysted animal; (C) schematic showing a sagittal section of the head; (D) Schematic of the position of cephalic neural structures; (E, F) SBEM images of a 6-month old cyst showing the elements forming the circumbuccal ring-shaped structure (star) innervating the mouth region; (G) 3D visualisation of the ventrolateral view of the cephalic neural structures from the same specimen as in E, F. Note that the dotted line shows the presence and altered morphology of the circumbuccal ring-shaped structure. Anterolateral cluster (ALc); circumoral sensory field (cosf); buccal crown (cr); retraction of the mouth area (mi); mouth opening (mo); ventral cluster (Vc); ventrolateral sensory field (vlsf); slit through which the mouth area retracts (asterisk).

The dorsolateral cluster was an unpaired neural cluster. The posterior part of this cluster formed three distinctive lobes – two lateral (left, right) and one median lobe. Anteriorly, the neurites that originated from the DLc formed the anterolateral sensory field (alsf) on both sides (left and right, respectively). The DLc extended laterally on both sides. The DLc’s lateral portions (left and right) were connected to the posterolateral cluster by a bridge (left and right, respectively). The posterolateral cluster (PLc) was paired, and the left and right PLc could be distinguished. The posterolateral cluster, an elongated cluster extending dorsoventrally, was the most posteriorly located relative to other neural clusters and the buccal-pharyngeal apparatus. Left and right PLc in the dorsal part formed the posterolateral sensory field (plsf, left and right, respectively). Ventrally, both posterolateral nervous clusters were connected with the ventrolateral sensory field (vlsf) on the corresponding side of the cephalic region (left or right). The integument covered all sensory fields (Figure 1a–f and 2a– d, f–h, Video S1). Under the buccal tube, the ventral cluster (Vc) was located on the ventral side of the head. The left and right anterolateral clusters (L-ALc and R-ALc, respectively) were noted on both sides of this unpaired cluster. In contrast to the PLcs, the ALcs were more extended anteriorly. Vc and both ALcs formed the ventrolateral complex (VLc) (Figure 1a,b,h,i, Video S1). The VLc and DLc elements were linked by somata regions (Figure 1g). The ventrolateral complex formed protruding extensions on both sides, closing in the upper part. Therefore, a circumbuccal ring-shaped nervous structure formed, and ventral and lateral portions of the ventrolateral complex co-created this ring-shaped structure. Extended nerves innervated the cuticle around the mouth opening, forming a circumoral sensory field (cosf) (Figures 2a,b,e,i, S2 and 3, Video S1).

### 3.2. Outer and inner connectives

Within the central nervous system, the cephalic neural clusters were connected to the rest of the central nervous system by two pairs of thick connectives. Each inner connective linked the VLc with the first trunk ganglion. The outer connectives connected the left and right posterolateral cluster with the same trunk ganglion as the inner connectives (Figure 1m, Video S2). The external lamina also surrounded the connectives.

### 3.3. Trunk nervous system

The ganglionated nervous system extended along the anteroposterior axis and showed a metameric pattern in which four large ganglia were positioned ventrally (Figure 4, Videos S2 and S3). A pair of thick interganglionic connectives (left and right) ran longitudinally from each ganglion and linked with the next ganglion up to the fourth one. The serial sections made it possible to find and visualise thin interpedal commissures. The three interpedal commissures in the trunk nervous system ran transversely to the longitudinally running connectives. Each of them was located between the two next ganglia. The first commissure was situated between the first and the second ganglia. The second commissure was present between the second and the third ganglion, whereas the last commissure was between the third and the fourth ganglion (Figure 4a–c, Videos S2 and S3). The structure of each trunk ganglion, similarly to the cephalic nervous clusters, was composed of the nerve fibre regions and somata regions rich in nerve cell bodies. Within the ganglia, synapses were observed. In contrast to the ganglia, soma was not found within the connectives and commissures. The elements of the ventral nervous system were externally surrounded by the external lamina (Figures 4e,h–l and S3).

**Figure 4.**
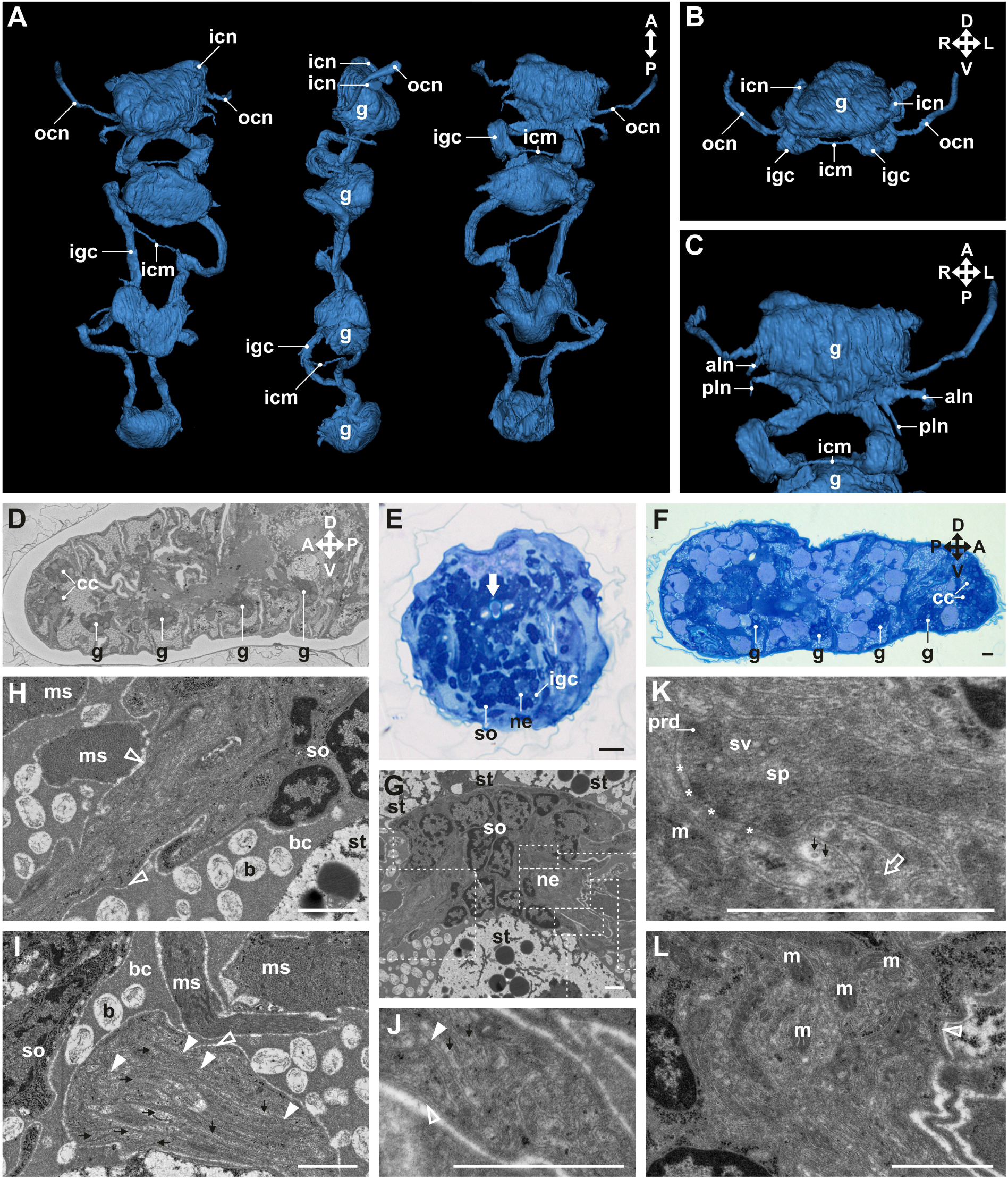
The central nervous system of the trunk in an encysted *Thu*. *ruffoi*. (A–C) 3D visualisation of the ventral neural structures of a 6-month-old cyst with SBEM image (D) from this specimen; (E) animal at an unknown time of encystment duration; (F) 8 to 9-month-old cyst; (G–L) freshly formed cyst; (E, F) LM, scale bar 10 µm; (G–L) STEM, scale bar 1 µm. Note that: A↔P indicates the anteroposterior axis, D↔V is the dorsoventral axis, while L↔R denote the left and right sides. Anterior leg nerve (aln); bacteria (b); body cavity (bc); cephalic neural cluster (cc); ventral ganglion (g); interpedal commisure (icm); inner connective (icn); interganglionic connective (igc); mitochondrion (m); somatic muscle (ms); neurites (ne); outer connective (ocn); posterior leg nerve (pln); presynaptic density (prd); somata (so); synapse (sp); storage cell (st); synaptic vesicles (sv); synaptic cleft (asterisk); basal lamina (empty arrowheads); interspaces between neural cells (filled arrowheads); bound dense material (empty thick arrow); buccal-pharyngeal apparatus (filled thick arrow); neurotubules (thin arrow).

Leg-specific innervation included two pairs of nerves extending from each side of the trunk ganglia. One of them was located more anteriorly (anterior leg nerve) than the other situated behind it (posterior leg nerve) (Figure 1c, Videos S2 and S3). Among the leg-specific neural structures, the leg ganglia were also observed, in which several nuclei were present. Within the fourth trunk ganglion, two pairs of nerves innervated the legs, (anterior and posterior leg nerves, respectively) and other structures, including two small peripheral ganglia on both sides of the animal in the posterior part of the contracted body. All leg-specific neural structures (leg nerves, leg ganglia) were not located within the legs of the encysted tardigrade but within the body cavity (Figure S3). The organisation of the trunk nervous system showed bilateral symmetry (Videos S2 and S3). Another pair of nerves was noted in the second and third trunk ganglia in front of the anterior leg nerve, one on each side of the ganglion. The second trunk ganglion innervated stomatogastric ganglion located close to the oesophagus and the anterior part of the midgut (Figure S3).

### 3.4. Changes during the cyst formation

The transition from the active (non-encysted) form to the immobile cyst involved changing the body’s general shape and morphology of the cephalic and extracephalic neural structures (Figures 2, S2, 3 and 5, Videos S1, S2 and S3). The general appearance of the cephalic nervous system organisation was compact. The cephalic region showed a tilt towards the ventral side that visibly changed the position of some structures, such as the dorsolateral cluster, which was located then much more frontally than dorsally. Also, posterolateral sensory fields were located much more anteriorly. On the opposite side, the ventrolateral sensory fields were directed towards the first trunk ganglion (Figures 2a–e, 3a–d and 5a–d, Videos S1 and S2). During cyst formation, the mouth area retracted inwards, which pressed the ring-shaped structure toward the neural cluster and distorted its morphology (Figures 2, S2 and 3, Video S1). The head was retracted into the rest of the body by bending the inner and outer connectives and narrowing the space between the ventrolateral complex and the first ventral ganglion of the trunk. These changes also resulted in the ventrolateral sensory fields being closer to the first trunk ganglion (Figure S5a–d, Video S2). Despite the symmetrical organisation of the cephalic neural structures, there were visible morphological changes in the head (Figure 1a–d, Video S1). Four ventral ganglia were connected by bent interganglionic connectives on both sides, which shortened the distance between the ventral ganglia. (Figures 4a,c and 5e–h). Moreover, morphological deformation was reflected by flattening the dorsal side of the second and third trunk ventral ganglia (Figure 4a, Videos S2 and S3). Additionally, changes within the legs during cyst formation resulted in the localisation of leg-specific structures within the body cavity (Figure S3).

**Figure 5.**
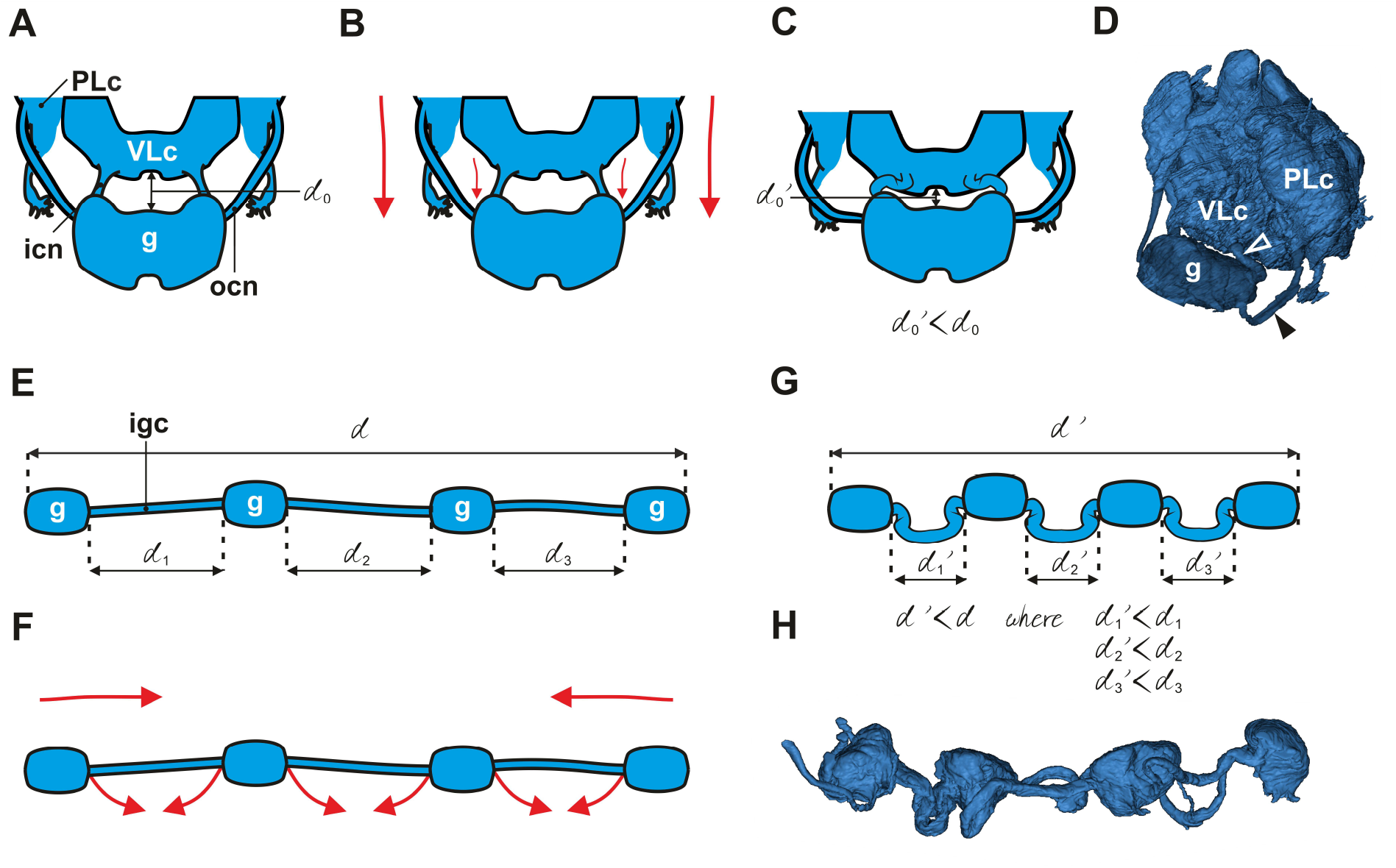
Changes within the nervous system during body contraction in cyst formation in *Thu*. *ruffoi*. (A–D) changes on the border between the head and trunk; (E–H) changes within the ventral nervous system of the trunk; (A, B, E, F) non-encysted animal; (C, D, G, H) encysted animal; (B) schematic of changes (red arrows) in which contraction of the body in the central direction includes contraction of the cephalic region into the rest of the body during cyst formation; (C) reduction in the distance (d_0_’ < d_0_) between the ventrolateral complex and the first ventral ganglion of the trunk. Note also the bending of the inner and outer connectives; (D) 3D visualisation of the link between the cephalic neural structures and the first ventral ganglion in a 6-month-old cyst; (F) schematic of changes (arrow) during which cyst formation involves contraction of the whole body in the central direction and changes within the ventral nervous system of the trunk; (G) reduction in the distance between the ventral ganglions (d_1_’ < d_1_, d_2_’ < d_2_, d_3_’ < d_3_) results in general distance reduction (d’ < d) between the two distal ends of the ventral nervous system of the trunk. Note also the bending of the interganglionic connectives; (H) 3D visualisation of the ventral nervous system morphology in a 6-month-old cyst. Ventral ganglion (g); inner connective (icn); interganglionic connective (igc); outer connective (ocn); posterolateral nervous cluster (PLc); ventrolateral complex (VLc); altered morphology of the inner (empty arrowhead) and outer (filled arrowhead) connectives.

## 4. DISCUSSION

### 4.1. Organisation of the nervous system

Data on the internal organisation of encysted tardigrades are sparse, including the nervous system organisation. Elements of the central and peripheral nervous systems are present in encysted individuals of *Thu*. *ruffoi* up to the 11th month of encystment duration. Despite the necessity to use interpolation in the segmentation of the cephalic region, the data obtained provided information on the nervous system’s organisation in this encysted species. In this paper, we have used the terminology found in the literature to describe the different parts of the nervous system (Walz, 1978; Wiederhöft & Greven, 1996, 1999; Zantke et al., 2008; Biserova & Kuznetsova, 2012; Persson et al., 2012, 2014; Schulze & Schmidt-Rhaesa, 2013; Schulze et al., 2014; Gross & Mayer, 2015; Mayer et al., 2013a, b; Gross et al., 2021), applying these terms as consistently as possible and retaining the most descriptive characters. In other cases, new terms have been established, which we believe better reflect the description of specific structures. Although the presence of glial cells was described based on a high number of ribosomes in *Macrobiotus* cf. *hufelandi* (Greven & Kuhlmann, 1972), we believe, similar to Wiederhöft & Greven (1996), that the presence of numerous ribosomes is not sufficient to delineate glial cells.

#### Head and sensory organs

Different authors have used various terms describing the brain parts and have interpreted ’’the brain’’ in other ways (compare, e.g. Schulze & Schmidt-Rhaesa, 2013 and Gross et al., 2021). Also, the issue of brain segmentation issue has been disputed by various authors (Persson et al., 2012, 2014; Mayer et al., 2013a; Gross & Mayer, 2015; Smith et al., 2016, 2018). The brain of the encysted tardigrade *Thu. ruffoi* is symmetrical and relatively compact, with the dorsolateral cluster being a large, dorso-ventrally flattened cluster. Posteriorly, there are three lobes, as also reported in non-encysted *Milnesium* cf. *tardigradum* and *Macrobiotus* cf. *harmsworthi* (Marcus, 1929; Wiederhöft & Greven, 1996; Mayer et al., 2013a). The posterior-most cluster of the cephalic region in the encysted *Thu. ruffoi* is the posterolateral one (also known as the outer cluster or outer lobe *sensu,* e.g. Wiederhöft & Greven, 1996; Mayer et al., 2013b; Gross et al., 2021, or posterior cluster *sensu,* e.g. Schulze et al., 2014), which is present on both sides. The dorsolateral cluster is connected with each PLc by a structure called a bridge. Comparative analysis of this region clearly shows the presence of neurites that link the posterolateral cluster with the dorsolateral one (Schulze & Schmidt-Rhaesa, 2013; Gross et al., 2021).

An interesting but also controversial issue is the presence/absence and nature of neural tissue in the ventral/ventrolateral part of the head. Among eutardigrades, ventrolateral accumulation of the neural tissue was found in *Mil*. cf. *tardigradum* (Apochela) (Wiederhöft & Greven, 1996) and *Halobiotus crispae* (Parachela) (Persson et al., 2012). Surprisingly, a similar pattern of organisation is present in *Thu*. *ruffoi* but not found in other parachelans such as *Macrobiotus* sp. or *Hypsibius* sp. (Zantke et al., 2008; Mayer et al., 2013a, b; Gross & Mayer, 2015). Moreover, the connection between protruding extensions of the VLc and the dorsolateral cluster, and the participation of these extensions in the formation of the circumbuccal ring-shaped structure resembles conclusions gathered from analyses of *Mil*. cf. *tardigradum* (Wiederhöft & Greven, 1996).

The present study shows that in encysted *Thu*. *ruffoi*, at least four types of sensory fields are associated with the integument: the circumoral sensory field, paired anterolateral sensory field, paired ventrolateral sensory field and paired posterolateral sensory field. The circumoral sensory field innervating the peribuccal region has been described in echiniscid (Heterogardigrada) and apochelan and parachelan (Eutardigrada) species (Walz, 1978; Wiederhöft & Greven, 1996, 1999; Dewel & Eibye-Jacobsen, 2006; Biserova & Kuznetsova, 2012). The dorsolateral cluster innervating the anterolateral sensory field on both sides has been well-studied by other researchers in different species of eutardigrades (Walz, 1978; Wiederhöft & Greven, 1996, 1999; Persson et al., 2012; Mayer et al., 2013a). This cluster is also associated with the presence of the median cirrus homolog found in Echiniscoidea (Dewel & Eibye-Jacobsen, 2006; Schulze & Schmidt-Rhaesa, 2013; Gross et al., 2021) and some Parachela (Zantke et al., 2008; Persson et al., 2012) representatives. Due to the lack of data, the presence of the median sensory field in *Thu*. *ruffoi* could not be verified, but its presence could not be excluded. Many researchers (e.g. Wiederhöft & Greven, 1996; Zantke et al., 2008; Persson et al., 2012; Mayer et al., 2013a) have indicated that the region of each posterolateral cluster is associated with the posterolateral sensory field (temporalia *sensu* Kristensen, 1982), which corresponds with data obtained for encysted *Thu*. *ruffoi*. What is even more interesting is that the posterolateral clusters also participate in innervating the ventrolateral sensory field in encysted *Thu*. *ruffoi*. Also, the foregut-associated sensory organs, such as the suboral region (also known as the suboral sensory fields) and buccal organ, are present among both parachelan and apochelan eutardigrades (Walz, 1978; Wiederhöft & Greven, 1996, 1999; Biserova & Kuznetsova, 2012) but were not studied in the present analysis. Moreover, they were also found in an echiniscid heterotardigrade (Dewel & Eibye-Jacobsen, 2006)

#### Head-trunk link

Both pairs of connectives (outer and inner) link the cephalic elements with the trunk nervous system. Results presented here show that the inner connectives link the first ventral trunk ganglion with the ventrolateral complex. Similar results were pointed out previously for heterotardigrade *Echiniscus testudo* (Echiniscoidea) (Schulze & Schmidt-Rhaesa, 2013; Gross et al., 2021) and eutardigrades *Mil*. cf. *tardigradum* (Wiederhöft & Greven, 1996) and *Hal. crispae* (Parachela) (Persson et al., 2012). Based on studies of *Mac.* cf. *hufelandi* and *Mac*. cf. *harmsworthi* (Zantke et al., 2008; Mayer et al., 2013a, b), the inner connective seems to pass near the ring-shaped structure and is linked to the dorsally located brain. Therefore, the inner connectives in *Thu*. *ruffoi* may run through the lateral part of the ventrolateral complex and link with the dorsolateral cluster. The entry place of outer connectives in *Ech*. *testudo* could not be localised by Schulze & Schmidt-Rhaesa (2013), whereas Gross et al. (2021) indicated that the outer connectives run directly into the dorsolateral cluster (brain *sensu* Gross et al., 2021) in this species. In contrast, data obtained for encysted *Thu*. *ruffoi* clearly show that the outer connectives enter the posterolateral clusters and then probably run through to the dorsolateral one, which seems to be confirmed based on data reported by Mayer et al. (2013a). Analysis of these data suggests that some fibres running from the outer connective within posterolateral clusters have a branch running to the posterolateral sensory field. Similar results have been found in *Mil*. cf. *tardigradum* (Wiederhöft & Greven, 1996), *Mac.* cf. *hufelandi* (Zantke et al., 2008) and *Hal. crispae* (Persson et al., 2012).

#### Trunk

The general pattern of the trunk nervous system organisation in encysted *Thu*. *ruffoi* corresponds to descriptions of other species (e.g. Mayer et al., 2013a; Schulze & Schmidt-Rhaesa, 2013; Schulze et al., 2014). The presence of synapses in encysted *Thu*. *ruffoi* confirms the recent findings (Gross et al., 2021) about synapsin-like rich regions within all of the trunk ganglia – in contrast to a previous report (Schulze et al., 2014), in which labelling with the antibody directed against synapsin was not positive for trunk ventral ganglia. Interestingly, the peripheral nerve of the second ventral ganglion innervates the stomatogastric ganglion in encysted *Thu*. *ruffoi*, as was described for non-encysted individuals of *Mac*. Cf *harmsworthi* (Mayer et al., 2013a, b). As in *Thu*. *ruffoi*, the leg ganglia and presence of the leg and peripheral nerves were reported for non-encysted eutardigrades, such as *Hyp. exemplaris* (Gross & Mayer, 2015), *Mac*. cf. *harmsworthi* (Mayer et al., 2013a, b) and *Hal. crispae* (Persson et al., 2012). These neural structures were also cited in heterotardigrades (Schulze et al., 2014; Schulze & Schmidt-Rhaesa, 2013; Gross et al., 2021). Additionally, a peripheral ganglion posteriorly located within the body of encysted *Thu*. *ruffoi* was also found, similar to the peripheral ganglion described by Mayer et al. (2013b) and Zantke et al. (2008, figure 5E).

### 4.2. Encystment and the nervous system

Histolysis of internal organs during encystment has been suggested (Murray, 1907a, b; Heinis, 1910; Richters, 1909; Hansen & Katholm, 2002). However, Von Wenck’s analysis of sectioned cysts did not support this hypothesis (von Wenck, 1914). Von Wenck (1914) studied *Mac. lacustris*, presently considered *Thu. augusti* (Degma & Guidetti, 2023), stating “[…] *it was clear that the* […], *eyes and brain were unchanged* […]” (von Wenck, 1914, p. 499). This statement may be considered evidence that the nervous system is unchanged by histolysis but is insufficient as it does not describe the organisation of the nervous system. Therefore, our data provide the first description of the nervous system’s organisation and morphology in an encysted tardigrade, a representative of the family Doryphoribiidae. Data collected in this study show that histolysis is not a part of encystment itself.

Changes in the morphology of the nervous system occur during cyst formation. Retraction of the mouth area and tilting of the head more ventrally occur during cyst formation. Cyst formation entails contraction of the whole body, compressing the head into the rest of the body and changing the specific morphology of the trunk nervous system. Compressing the head into the rest of the body and retracting the mouth compacts the organisation of the cephalic nervous system, which is similar to that seen in contracted non-encysted *Mac*. cf. *harmsworthi*. Similarities between encysted and non-encysted tardigrades are also present with respect to the morphology of the connectives (Mayer et al., 2013a). The entire trunk contracts, which can be assumed based reduction in distance between ventral trunk ganglia. Atypical morphology in the head or trunk that may occur during cyst formation is apparently related to the contraction of the somatic muscles of the individual and pressures from the body wall and the organs and cells in the body cavity. Since the leg-specific neural structures of all four pairs were located within the body cavity, their position is far from the natural position described in other non-encysted tardigrades (Gross & Mayer, 2015; Persson et al., 2012; Mayer et al., 2013a, b; Schulze & Schmidt-Rhaesa, 2013; Schulze et al., 2014; Gross et al., 2021).

### 4.3. Nervous system activity during encystment

The presence of the presynaptic and postsynaptic densities in encysted *Thu*. *ruffoi* supports the conclusion that some activity of the nervous system occurs during encystment. The electron-dense region of the presynaptic area (called “an active zone”) is associated with the site of synaptic vesicle fusion with the presynaptic membrane and exocytosis of neurotransmitters into the synaptic cleft (Emperador-Melero & Kaeser, 2020). In contrast, the postsynaptic area is an electron-dense, protein-rich structure beneath the postsynaptic membrane responsible, among other functions, for receiving and interpreting signals transmitted by presynaptic termini (Sheng & Kim, 2011; Cohen, 2013).

### 4.4. Future prospects

Further research on the nervous system organisation of *Thu*. *ruffoi* is warranted. Gaps in the data and the compact morphology of the cephalic neural structures in encysted animals justify that future research should focus on additional morphological analysis, especially of the anterior part and medial region, to fully determine the morphology of the cephalic nervous system of both encysted and non-encysted tardigrades. Additional analyses (e.g. immunohistochemistry) will further elucidate the cephalic nervous system organisation, especially the relationship between the ventrolateral complex and the dorsolateral cluster.

## 5. CONCLUSIONS

Based on the present study, the following conclusions have been reached: **1)** The nervous system of a freshwater eutardigrade during a period of 11^th^ months in encystment is composed of elements representing the central and peripheral nervous system; **2)** the circumoral, ventrolateral, anterolateral and posterolateral sensory fields are present in encysted animals and are associated with the integument; **3)** an unpaired stomatogastric ganglion innervates the oesophagus and the midgut; **4)** histolysis is not an integral part of the encystment process in this species; and **5)** cyst formation affects the animal’s morphology, including the nervous system, in a specified pattern.

## Authors’ contributions

K.J. – Conceptualisation, Data curation, Formal analysis, Funding acquisition, Investigation, Methodology, Validation, Software and Visualization, Writing (original draft preparation, review and editing); I.P. **–** Methodology, Validation, Writing (review and editing), Supervision

## Acknowledgements

We are grateful to the Michalczyk Lab (Jagiellonian University, Kraków, Poland), particularly Dr. Daniel Stec and Dr. Mateusz Sobczyk, for obtaining, identifying and providing the Thu.ruf_PL.014 strain. Moreover, we would also like to thank Dr. Danuta Urbańska-Jasik, Dr. Łukasz Chajec, and Dr. Izabela Potocka (University of Silesia in Katowice, Poland) for their technical assistance and Dr. Małgorzata Śliwińska (Laboratory of Imaging Tissue Structure and Function, Nencki Institute of Experimental Biology, Warszawa, Poland) for obtaining a set serial EM images. Moreover, we are deeply indebted to Dr. Diane R. Nelson for revision, valuable comments and improving the English style. We also thank the reviewers for any suggestions that helped to improve this manuscript.

## Data availability statement

The data that supports the findings of this study are available in the supplementary material of this article

