## Supplementary material for "Organisation of the nervous system in cysts of the freshwater tardigrade *Thulinius ruffoi* (Parachela, Isohypsibioidea: Doryphoribiidae)": Figure S

### Supplementary Figures

#### Figure S1

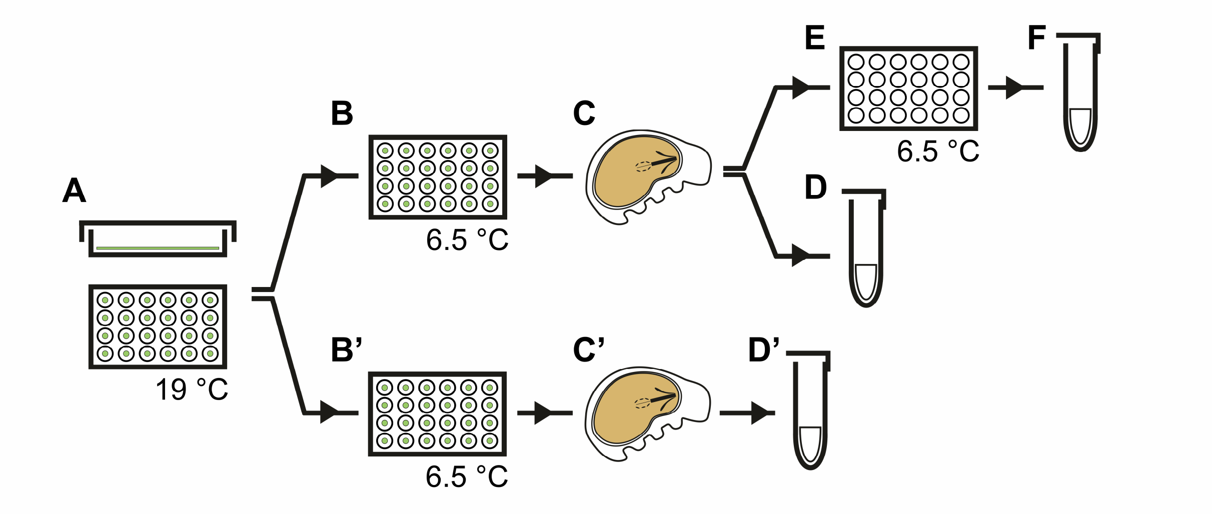


**Figure S1. Schematic for obtaining cysts of *Thu. ruffoi* for analysis.** (A) Non-encysted individuals from the control cultures in Petri dishes and 24-well plates (B, B') were placed (one animal per well) on the 24-well plastic plates and transferred to the lower temperature. (B–F) The animals were observed every 24–48 hours until cyst formation began to determine established encystment durations. After cyst formation (C), some freshly formed cysts (D) were isolated, cleaned and processed for analysis. Other freshly formed cysts (E) were isolated and put on a new plate to begin timing of cyst incubation. (F) After a specific incubation time, individuals that remained as cysts were then recovered, cleaned, and processed for analysis. (B'–D') Obtaining cysts with an indefinite duration of encystment was simpler. (B') The wells of the plates were observed occasionally (the min/max interval between checks was seven days and six months, respectively). (C') If cysts were present, some were (D') randomly selected, recovered, cleaned and processed for analysis. This activity could be repeated because the eggs and juveniles on the plates (B') were not removed, and the population grew. Note that the green colour indicates the presence of food (algae).

### Figure S2

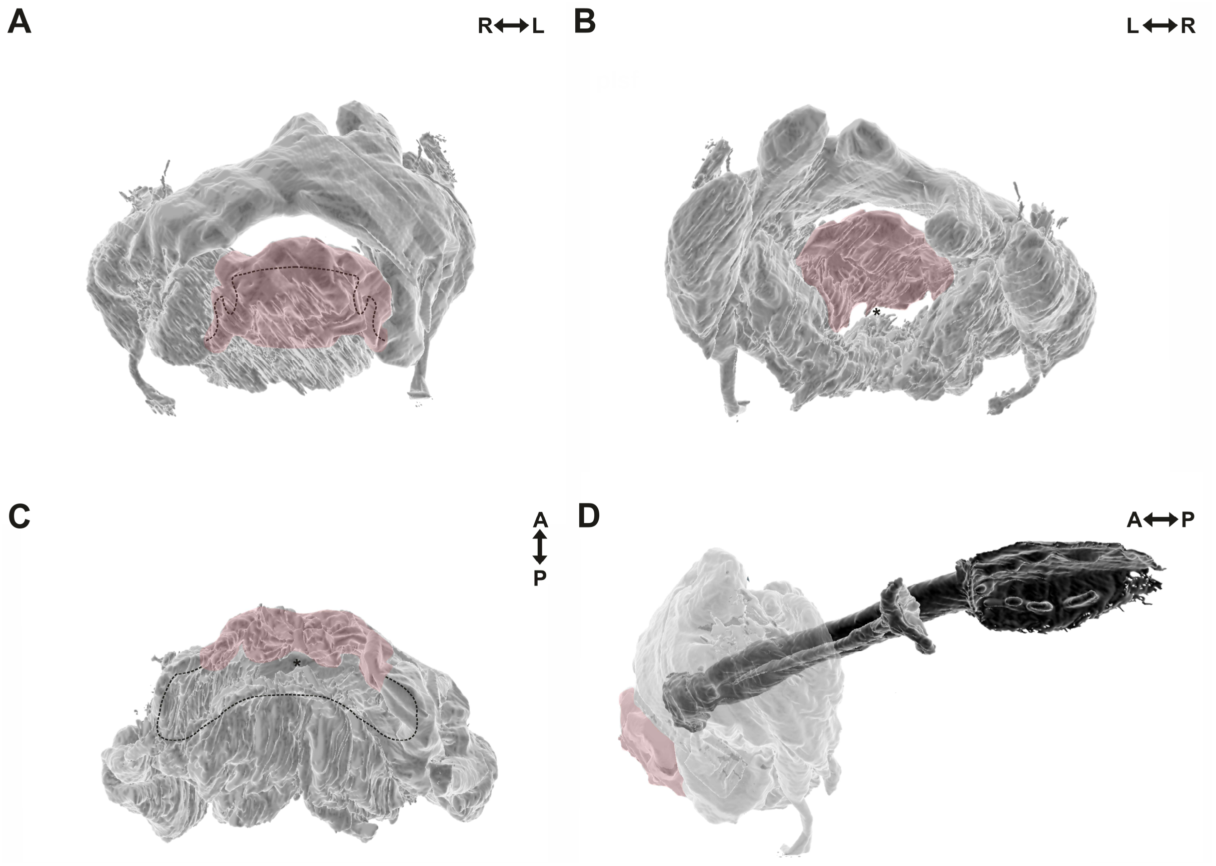


**Figure S2. Circumbuccal ring-shaped structure of *Thu. ruffoi* and its altered morphology.** (A) frontal view; (B) back view with the ring-shaped structure and a slit (in the centre) through which the innervated mouth area was retracted; (C) ventral view; (D) colocalisation of the externally extended part of the circumbuccal ring-shaped structure to the buccal-pharyngeal apparatus. A dotted line shows the appropriate part (frontal in A, or ventral in C) of the circumbuccal ring-shaped structure and its altered morphology. The retracted mouth area goes through a visible slit (asterisk) and is innervated by surrounding nerve ring nerves. A false-coloured (red) part indicates the visibly extended part of the nerve ring pressed into the rest of the cerebral nervous tissue. A↔P indicates the anteroposterior axis, while L and R denote the left and right sides.

### Figure S3

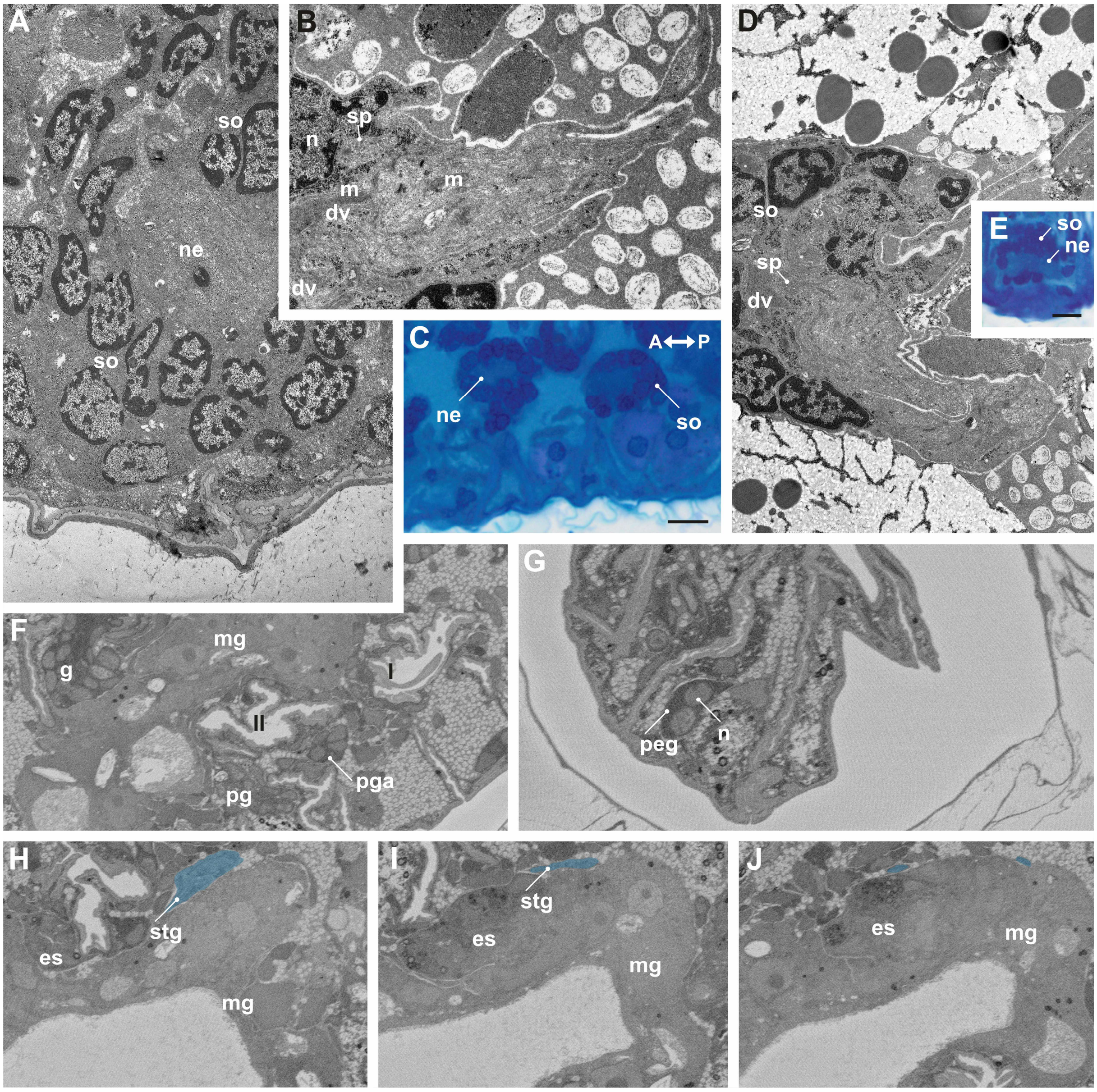


**Figure S3. Trunk central nervous system and peripheral neural structures of encysted *Thu. ruffoi*.** (A–E) ventral ganglia with fragments of interganglionic connectives (B, D); (F) leg ganglion innervated by second ventral ganglion; (G) peripheral ganglion innervated by fourth trunk ganglion; (H–J) stomatogastric ganglion innervated by the second ventral ganglion and its processes running in the oesophagus and midgut direction. Note that the Roman numerals indicate the appropriate pair of legs; (A, C, E) three different cysts with indefinite durations of encystment; (B, D) freshly formed cyst; (F–J) a 6-month-old cyst; (A, B, D) TEM, scale bars 1 µm; (C, E) LM, scale bars 10 µm; (F–J) SBEM images. Bound dense material (dv); oesophagus (es); ventral ganglion (g); mitochondrion (m); midgut (mg); nucleus (n); neurites (ne); peripheral ganglion (peg); pedal gland (pg); leg ganglion (pga); somata (so); synapse (sp) with synaptic vesicles; stomatogastric ganglion (stg).
