## Supplementary material for "Organisation of the nervous system in cysts of the freshwater tardigrade *Thulinius ruffoi* (Parachela, Isohypsibioidea: Doryphoribiidae)": Table S1

| | number of cysts | purpose |
|---|---|---|
| *cysts with a well-established time of encystment duration* | **19 in total** | |
| freshly formed (up to 48h) | 6 | TEM/STEM/LM |
| 1mth | 1 | TEM/STEM/LM |
| 3mths | 5 | TEM/STEM/LM |
| 6mths | 4 | TEM/STEM/LM (3);<br>SBEM & 3D visualisation (1) |
| 8-9mths | 2 | TEM/STEM/LM |
| 11mths | 1 | TEM/STEM/LM |
| *cysts without an established time of encystment duration* | **11 in total** | TEM/STEM/LM |
| | Obtained cysts: **30** | |

**Table S1. List of cysts of *Thu. ruffoi* which were obtained and used for specific purposes.** Parentheses indicate the number of individuals used in different analyses, if applicable.
