## Supplementary material for "Organisation of the nervous system in cysts of the freshwater tardigrade *Thulinius ruffoi* (Parachela, Isohypsibioidea: Doryphoribiidae)": Video S

### Supplementary Videos

**Video S1. The cephalic neural structures of encysted *Thu. ruffoi*.** Note the altered morphology and the location of the sensory fields.

**Video S2. Cephalic and trunk nervous system of encysted *Thu. ruffoi*.** Note the very visible connection between the ventral and cephalic part of the central nervous system by the inner and outer connectives. Peripheral structures of the trunk and leg neural structures are not shown.

**Video S3. The ventral nervous system of encysted *Thu. ruffoi*.** Note visibly changed morphology due to compression caused by body contraction. Peripheral structures of the trunk and leg neural structures are not shown.
